# Nanopipettes enable native electrospray ionisation ion mobility mass spectrometry studies of buffer-dependent conformational dynamics of α-synuclein

**DOI:** 10.1101/2025.08.19.671163

**Authors:** Emily J. Byrd, Emma L. Norgate, Joel A. Crossley, Chalmers C. C. Chau, Bob Schiffrin, Alexander Kulak, Sheena E. Radford, Paolo Actis, Antonio N. Calabrese, Frank Sobott

## Abstract

Electrolyte conditions *in vivo* and *in vitro* are known to influence protein structure and function. Intrinsically disordered proteins (IDPs) are particularly sensitive to their solution conditions such as ionic strength and molecular crowding, and their dynamic structural ensembles rapidly respond to the cellular environment. While structural mass spectrometry (MS) techniques are uniquely able to capture aspects of this structural diversity, technical limitations have largely precluded the use of native MS approaches to interrogate the conformational rearrangements of IDPs in response to high concentrations of non-volatile salts. Here, we overcome this challenge by employing sub 100-nm nanopipette electrospray emitters for more gentle and salt-tolerant analysis to study the conformational distribution of α-Synuclein (αS) using native MS and ion mobility-MS in varied solution conditions, including in phosphate buffered saline. We show using native MS that it is possible to capture salt and buffer induced changes in the αS conformational ensemble when using traditional biochemical buffers, which reflect structural changes from *in silico* predictions and in-solution measurements. This work demonstrates the power of nanopipette emitters for the study of IDPs, and establishes native MS as a method that can be routinely used to determine how solution conditions tune the conformational landscape of IDPs.

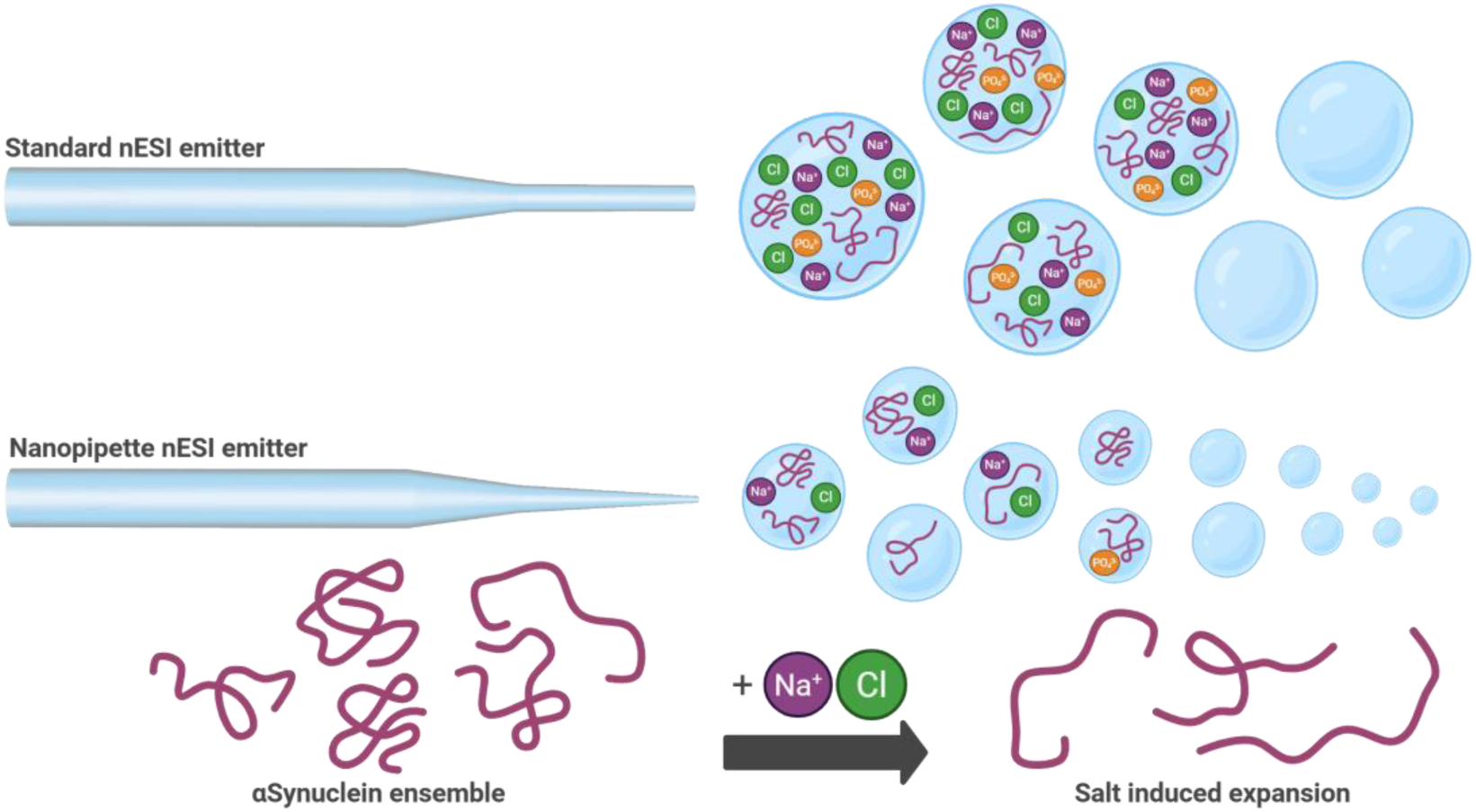

## INTRODUCTION

Intrinsically disordered proteins (IDPs) and proteins containing intrinsically disordered regions (IDRs) partake in many protein-protein interactions, aided by their structural plasticity, which facilitates their functional roles in processes such as transcriptional regulation and cell signalling^[1–4]^. IDPs are known to explore a vast landscape of rapidly inter-converting and co-existing configurations which are determined by their sequence and the chemical composition of their environment^[5]^. Many IDPs and IDRs are implicated in disease pathogenesis and are therefore therapeutic targets. This includes diseases such as cancer^[6,7]^ and neurodegenerative conditions, where many IDPs have been shown to undergo a disorder-to-order transition to form insoluble, β-sheet rich amyloid fibrils ^[8–10]^. Deposits of amyloid fibrils are associated with a number of neurodegenerative diseases, including Parkinson’s disease (PD), where α-synuclein (αS) fibrils are found in insoluble Lewy body deposits in the substantia nigra of the brain ^[11–17]^.

The structure and function of the IDP αS are affected by environmental conditions. For example, increasing NaCl concentration (from 50 mM up to 1000 mM) enhances liquid-liquid phase separation of αS in the presence of crowding agents (PEG400)^[18]^, by exposing the non amyloid-β component (NAC) domain, a hydrophobic region of the sequence (residues 60-95) through the displacement of water molecules^[19]^. This ultimately results in modulation of the aggregation pathway, leading to different fibril morphologies^[20–23]^. Moderate physiological concentrations of NaCl (up to *ca.* 0.1-0.3 M) have been shown to weaken contacts between the N-terminal, C-terminal and the NAC domain of αS, resulting in fewer intramolecular contacts and enhancing aggregation into amyloid^[24,25]^. Additionally, monovalent and divalent ions such as Ca^2+^, Cu^2+^, Cu^+^, Mn^2+^, Mg^2+^, Fe^2+^ and Zn^2+^, many of which are shown to be present at elevated concentration in the brains of Parkinson’s disease patients and are colocalised with αS in Lewy body assemblies^[26,27]^, have been identified as regulators of the conformational landscape of αS and as accelerators of amyloid assembly^[26,28–38]^. Previous data from hydrogen-deuterium exchange-mass spectrometry identified that the conformational behaviour of αS differs between solution conditions that match the extracellular, cytosolic and lysosomal environments (which differ in pH and ionic strength), with discrete conformational changes identified in the central hydrophobic NAC domain of αS which forms the amyloid core^[39,40]^. These different solution conditions also result in altered αS amyloid assembly kinetics^[39,41,42]^, highlighting the importance of replicating physiological conditions when studying IDPs *in vitro*.

Native mass spectrometry (native MS), especially when combined with ion mobility-MS (IM-MS), is able to capture and characterise dynamic, transient, heterogeneous species within a conformational ensemble and is a powerful technique for investigating IDP conformational ensembles^[43–50]^. The first step in a native MS experiment is the gentle ionisation and transfer of protein species which are kinetically trapped in their solution states into the gas phase. This process is typically achieved using nano electrospray ionisation (nESI) under non-denaturing solution conditions so that information about protein conformations adopted in solution can be measured in the gas phase^[51,52]^. The desolvation process that occurs during nESI requires the use of aqueous solutions comprising volatile salts such as ammonium acetate (AmAc)^[53]^. The presence of salts that are commonly required during protein preparation (such as Na^+^, K^+^, PO_4_^3+^ and Tris) as well as buffer additives that may be critical to maintain the structural integrity of proteins, can cause ion suppression which reduces spectral quality, resolution and sensitivity ^[54–57]^. Salt ions have also been shown to adduct to protein ions^[58–60]^, which can result in unresolvable mass spectra. Therefore, prior to native MS analysis, proteins are typically buffer exchanged from their biochemical storage buffer (a process which has been simplified recently with online buffer exchange^[61,62]^) into AmAc to dilute/remove salt ions from solution to prevent adduction in mass spectra (ammonium ions are volatile and do not adduct to proteins). However, many proteins require salts to remain stable in solution, and produce poorly resolved native mass spectra in AmAc^[60]^. One way to circumvent ion adduction is to generate smaller droplets during the nESI process so that salt ions remain independent from the solvent droplets containing protein molecules^[63–65]^. This approach has been applied successfully to the study of membrane proteins in detergents and other proteins in biochemical buffers^[59,66]^. Using IM to measure the rotationally averaged collision cross section (CCS) distributions of ion species coupled to MS isa powerful technique for studying the conformational properties of IDPs ^[35,36,50,67–70]^, and we and others have previously used this approach to probe the conformations of αS under different solution conditions (e.g. in the presence of metal ions, dopamine and detergents)^[35,36,38,50,71–74]^. However, these experiments have been limited to using AmAc based solutions.

Here we demonstrate the application of quartz nanopipette nESI emitters with a tip diameter of *<* 100 nm (standard borosilicate nESI emitters have diameters of *ca.* 2-20 µm; Figure S1 and S2) for native MS of an IDP, αS in its N-terminally acetylated form^[53,75,76]^. While previous work has shown that submicron emitters can be used for nESI and that they enable the acquisition of native mass spectra in diverse buffer conditions^[57,63,65,66,77,78]^, these efforts were limited to proof-of-concept studies of stably folded test proteins. Nanopipette emitters have not been previously used to elucidate structural changes involving αS, or other disease relevant IDPs. Moreover, the use of nanopipettes to characterise IDPs, a class of proteins that are especially responsive to the solution conditions^[67,79,80]^, and the conformational changes that they undertake, has not been previously described. Given the role of solution conditions in tuning the kinetics of αS aggregation^[34,39,40,42]^ and the experimental evidence demonstrating that different salt (NaCl) concentrations alter structural properties^[20,24,42]^, we chose to interrogate how ionic strength and buffer composition, known modulators of αS conformation and amyloid propensity^[18–20,42,81]^, tune conformational distribution, as measured by IM-MS. We show that native mass spectra of αS can be acquired in nonvolatile buffers commonly used in biochemical/biophysical studies of protein structure and function *in vitro*, including Dulbecco’s phosphate buffered saline (PBS), Tris-HCl, potassium phosphate and sodium phosphate, and in the presence of added NaCl. By combining data from IM-MS with *in silico* and in solution measurements of protein conformation, we demonstrate that native MS can reveal how both buffer composition and ionic strength tune the conformational heterogeneity of αS. Combined, this work demonstrates that nanopipette nESI emitters can expand the repertoire of possibilities for native MS and IM-MS studies of IDPs, to enable new insights into how IDP conformational ensembles respond to changes in the solution environment and understand how this impacts biological function or in the case of disease, dysfunction.

## RESULTS AND DISCUSION

### Native MS in physiological buffers

To test the limits of nanopipette technology for nESI-MS analysis of αS, we first attempted to use nanopipette emitters (quartz nanopipettes with a nanopore opening of *ca.* 40 nm, Figure S1 and Figure S2) to obtain native mass spectra of αS in PBS (Dulbeccos A), a buffer commonly used in biochemical studies. Excitingly, we were able to measure a well-resolved spectrum when using these nanopipette emitters, but (as expected) not when using standard µM-wide emitters (Figure 1a,b). A similar αS charge state distribution was observed with both emitters, suggesting that the smaller tip diameter of the nanopipette does not alter the structural ensemble of monomeric αS, as the charge states adopted in native MS relatively reflects a proteins solvent exposed surface area^[52,82,85–87]^.

**Figure 1.**
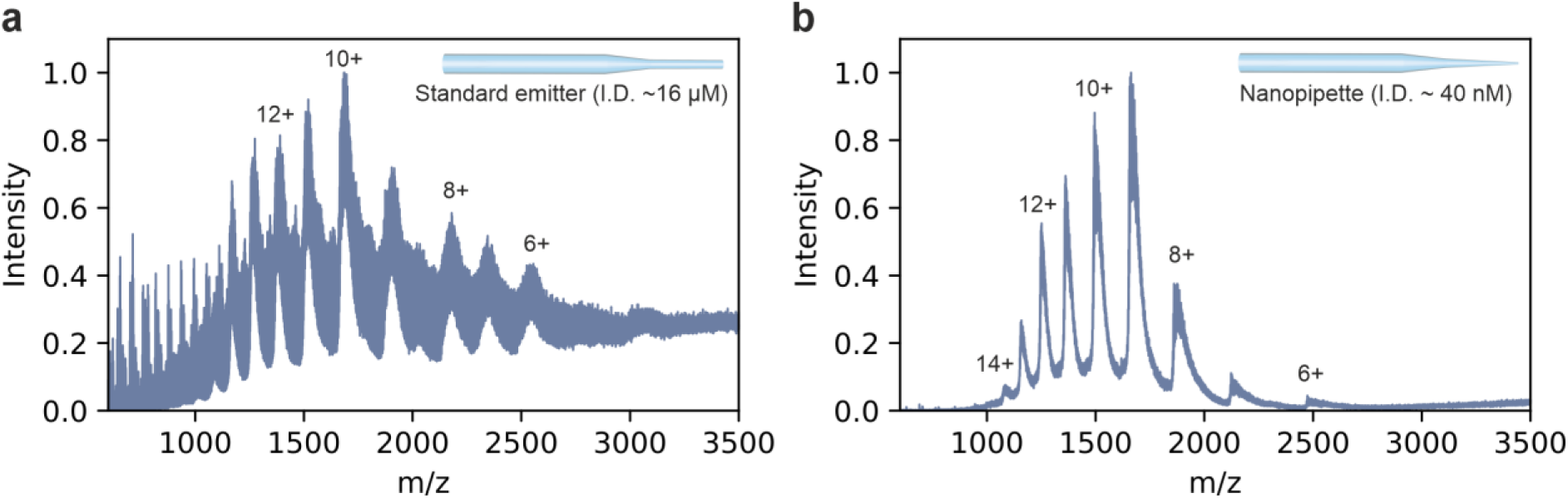
Direct comparison of nanopipette and standard nESI emitters. Representative native nESI mass spectra of 20 µM αS analysed in PBS, pH 7.2 using (a) standard emitters and (b) nanopipette emitters using identical instrument acquisition parameters on Waters Synapt HDMS (see Methods). I.D.: internal diameter.

In native MS analysis it is common to observe non-specific adducts and clusters of salts, e.g. Na^+^ (+23 Da) and K^+^ (+39 Da)^[83]^. Reduced salt clustering and salt adducts can be observed with nanopipette emitters (Figure 1b) compared to standard emitters (Figure 1a and Figure S3), consistent with previous reports^[54]^. The native mass spectrum in Figure 1b shows an overall decrease in salt adducts,as well as a marked reduction of salt clusters < 1000 *m/*z. Overall, this shows that nanopipette emitters enable well-resolved nESI mass spectra of αS in biochemical buffers, consistent with previous studies demonstrating an enhanced salt tolerance^[57,84]^. By showing that it is possible to capture the conformational diversity of αS in PBS using MS, we highlight the potential of nESI with nanopipette emitters for conformational analysis of IDPs in complex solution environments which mimic more biologically relevant conditions (cellular electrolytes, salts, crowding and liquid-liquid phase separation promoting agents), and demonstrate that there is no need to avoid non-volatile salts in native IM-MS.

When comparing native nESI mass spectra of αS acquired using nanopipettes in PBS (Figure 1b) with those in AmAc buffer (Figure 2a), a shift in the charge state distribution (6+ to 14+ vs. 5+ to 18+) can be noticed as well as the emergence of dimer spectral peaks in AmAc. The most intense charge state in PBS is 9+ whereas the 7+ charge state was most intense in AmAc, indicating a modulating effect of salt on αS conformational behaviour.

**Figure 2.**
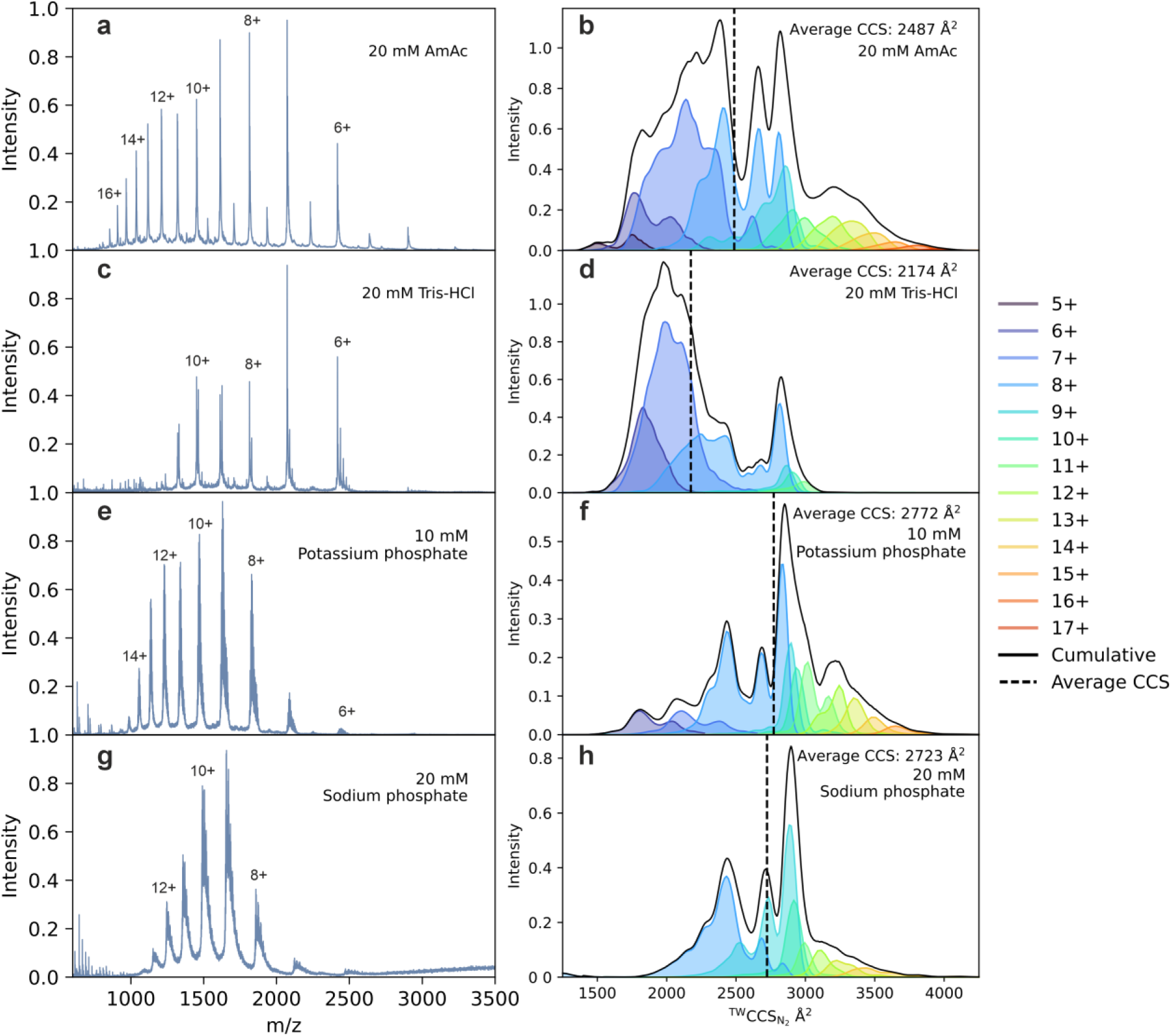
Native MS and IM-MS of αS in biochemical buffers. Native nESI mass spectra of 20 µM αS (left panel) and ^TW^CCS_N2_ distributions (right panel) in (a,b) 20 mM AmAc, (c,d) 20 mM Tris-HCl, (e,f) 10 mM potassium phosphate, (g,h) 20 mM sodium phosphate. Selected charge states are labelled in the mass spectra. The key on the righthand side indicates the colour of the CCS distribution for each charge state, the black solid line is the cumulative fit and the black dotted line represents the average ^TW^CCS_N2_ value from the distribution.

We next performed native MS and IM-MS to further understand how these changes in the mass spectra are related to structural alterations of αS in a range of commonly used biochemical buffers, by calculating collision cross section (CCS) distributions from the IM-MS data (see Methods)^[46]^. Travelling wave (TW) IM measurements were performed in N_2_ buffer gas for each charge state. Consistent with community standards^[48]^ we refer to these values as ^TW^CCS_N2_. To provide a snapshot of the conformational ensemble captured by the native IM mass specta, we measured ^TW^CCS_N2_ values for each detected charge state and plotted a cumulative distribution where the contributions from each individual charge state are intensity-weighted and combined (using a scaling factor based on the intensity of each charge state in the corresponding mass spectrum, see Methods). We acquired native mass spectra of αS from the buffers Tris-HCl, potassium phosphate and sodium phosphate, Figure 2c,e,g and measured IM-MS ^TW^CCS_N2_ distributions from these spectra (Figure 2b, d, f, h), for comparison with data acquired in conventional AmAc (Figure 2a,b)^[88,89]^.

Using nanopipette emitters, a resolvable charge state distribution was achieved for αS in each of the tested buffer systems, and clear shifts in charge state intensities were observed between buffers (Figure 2a,c,e,g). The native mass spectrum of αS in 20 mM AmAc (Figure 2a) appears as a multimodal distribution of charge states ranging from an extended conformational family (+17 to +10) to a compact conformational family (+9 to +5). This is reflected in the ion mobility ^TW^CCS_N2_ plot of these data (Figure 2b) and has been shown previously^[36]^. To enable a simple comparison between experimental conditions, we calculated the average (mean) CCS value from the culmulative ^TW^CCS_N2_ distributions (vertical dashed lines in Figure 2). In 20 mM AmAc, this average ^TW^CCS_N2_ value (2487 Å^2^) lies in between the extended and compact conformational families (Figure 2b). Interestingly, the spectra and ^TW^CCS_N2_ distributions in PBS (Figure 1b, Figure S4) and the other two phosphate buffers tested (Figure 2e-h) are comparable. All three phosphate buffered solutions resulted in ^TW^CCS_N2_ distributions that were narrower and favoured more expanded conformations compared to those observed in AmAc (average ^TW^CCS_N2_ values of 2598 Å^2^, 2772 Å^2^ and 2723 Å^2^ in PBS, potassium phosphate and sodium phosphate, respectively, compared to 2487 Å^2^ in AmAc). Contrary to this, the spectrum obtained in 20 mM Tris-HCl buffer is shifted to populate lower charge states, and charge state 12+ and above are not observed (Figure 2c). This is reflected in the ^TW^CCS_N2_ distribution which demonstrates that the compact conformational familes are favoured in Tris-HCl, with the average value of this ^TW^CCS_N2_ distribution being shifted to a lower value (2147 Å^2^; Figure 2d). It appears that in Tris-HCl, the conformational ensemble is less heterogeneous overall, populating two distinct conformational families, compared with a broader, multimodal distribution detected in AmAc, reflected by the decrease in the abundance of highly extended conformational families (higher charge states 12+ to 18+), however the ^TW^CCS_N2_ window in Å^2^ is the same. Conversely, the native mass spectra and ^TW^CCS_N2_ distributions of αS in 10 mM potassium phosphate, 20 mM sodium phosphate and PBS are consistent with more expanded αS conformational families predominating than in AmAc, and the CCS fingerprint appears distinctly different to αS in both AmAc and Tris-HCl.

Combined, these results show that the nm diameter orifice of the nanopipettes described in this work can enable acquisition of resolvable mass spectra to obtain IM-MS fingerprints of αS in a range of non-volatile biochemical buffers. The charge state distributions of αS observed in the mass spectra and the ^TW^CCS_N2_ distributions detected using IM-MS also differ in different buffer conditions. This suggests that modulation of the conformational landscape of the IDP αS in different buffers can be detected by native IM-MS using nanopipettes. Such a change has been reported previously for the folded protein bovine serum albumin (BSA) by analysis of native MS charge state distributions and gas phase thermal denaturation. AmAc was shown to stabilise compact conformational families of BSA and the presence of NaCl shifted the population of BSA towards more unfolded conformations^[84]^.

### Salt affects αS conformational behaviour

Next, we titrated NaCl into AmAc and Tris-HCl buffers to systematically investigate the effect of ionic strength on the conformational families populated by αS using IM-MS (Figure 3). In both AmAc and Tris-HCl, increasing the concentration of NaCl results in αS populating more highly charged species by enhancing the population of more expanded conformational families (Figure S5 and Figure S6, respectively with the 8+ charge state shown in Figure S7). This is supported by analysis of the ^TW^CCS_N2_ distributions which also show a clear shift towards higher average CCS values with increasing salt concentrations (Figure 3a-h). We hypothesise that this shift towards extended conformations is likely due to a charge screening effect in which Na^+^ ions interact with the negatively charged C-terminal region and Cl^−^ ions interact with the overall positively charged N-terminal region^[24]^, reducing intraprotein contacts. This is consistent with NaCl-dependent expansion of αS observed previously using small angle X-ray scattering (SAXS)^[19]^.

**Figure 3.**
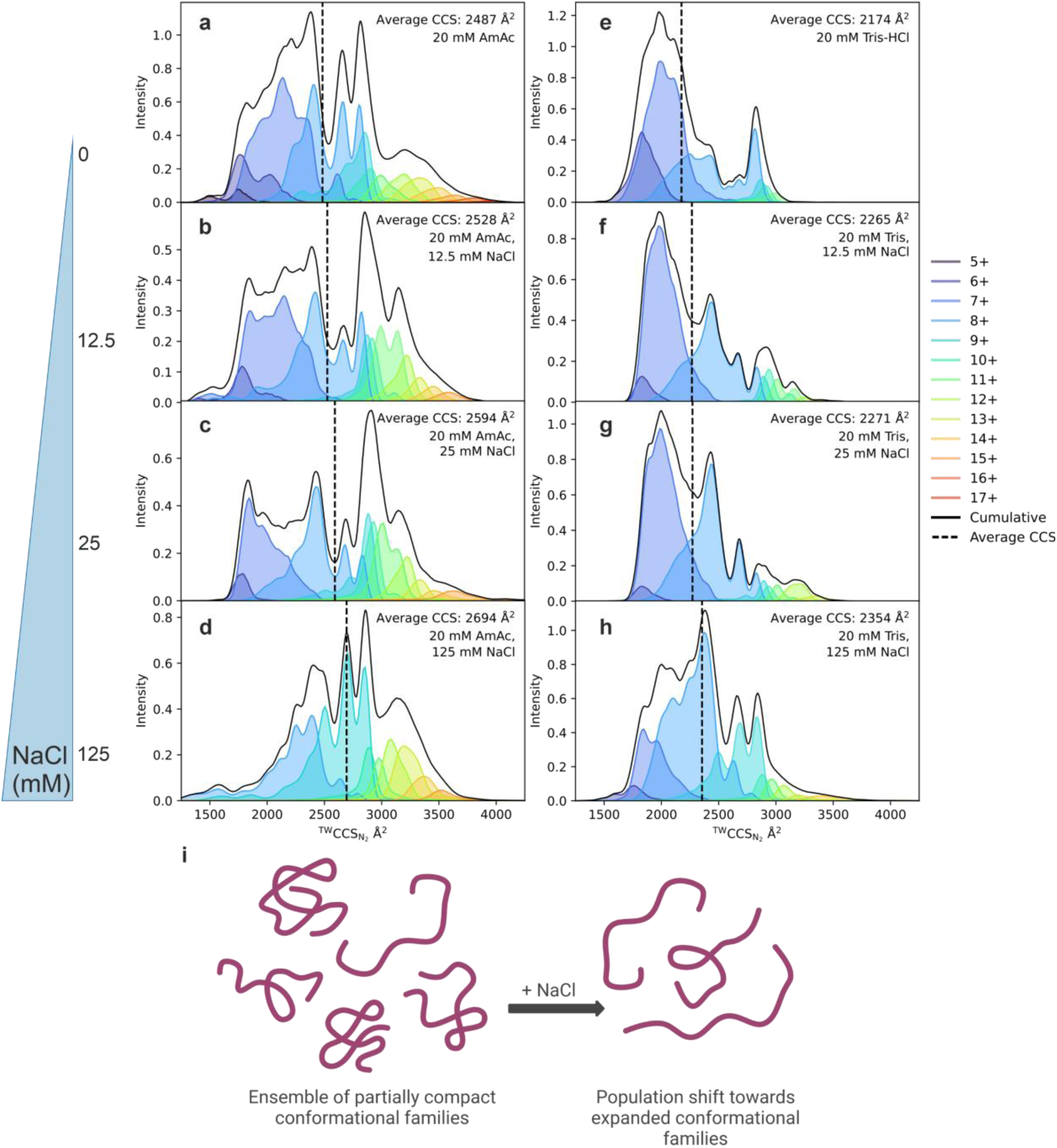
Salt titration of αS in AmAc and Tris-HCl by native IM-MS. ^TW^CCS_N2_ distributions showing the effect of salt (NaCl). αS was measured in (a) 20 mM AmAc with the addition of (b) 12.5 mM, (c) 25 mM and (d) 125 mM NaCl and also in (e) 20 mM Tris-HCl with the addition of (f) 12.5 mM, (g) 25 mM and (h) 125 mM NaCl. (i) A schematic of αS expansion in the presence of NaCl. The key on the righthand side indicates the colour identifying each charge state, the black solid line is the cumulative fit and the black dotted line represents the average ^TW^CCS_N2_ value.

As a control, we compared the effect of adding NaCl on the measured ^TW^CCS_N2_ values of αS with those of natively folded and disulfide-stabilised hen egg white lysozyme (Figure S8), a well-studied globular protein of similar molecular weight to αS (14,350 Da compared with 14,502 Da for N-terminally acetylated αS used here)^[90–92]^. In both AmAc and Tris-HCl buffers, more compact species and narrower conformational distributions of lysozyme were observed upon addition of 25 mM NaCl, by contrast with the conformational extension observed with αS. This suggests that our observations for αS are a result of the intrinsically disordered protein structurally changing in response to its solution environment, and that these structural features are preserved in the gas-phase and captured with IM-MS, and not a result of changes in the nESI process induced by the addition of salt. The results are consistent with a model in which the disordered and extended conformational families of αS are stabilised in elevated salt concetrations and that the protein is less driven to form intraprotein contacts which would stabilise more compact states (Figure 3i).

### Course grained molecular dynamics (MD) simulations suggests expanded states of αS are favoured at elevated NaCl concentrations

Previous coarse-grained MD simulations showed a salt dependent increase in the radius of gyration (R_g_) of αS up to a NaCl concentration of 0.1 M as a result of charge shielding^[24]^. We sought to expand on these studies using coarse-grained simulations of αS in implicit solvent using CALVADOS^[93,94]^. Simulations were carried out at ionic strengths ranging from 10 - 150 mM (these ionic strengths compare to the total ionic strength of 20 mM AmAc in the absence and presence of NaCl used in IM-MS experiments here) and the R_g_ values of the resultant model structures of αS were calculated. These data demonstrate a salt dependent increase in the average R_g_ of αS monomers from the simulations over the range of ionic strength values tested (Figure 4a). This change in average R_g_ is a result of shifts in the distribution of the αS conformational ensemble towards structures with higher R_g_ values with increasing ionic strength (Figure S9). We compared these computationally-generated R_g_ values with the apparent R_g_ (R_app_) derived from the weight-averaged mean of the ^TW^CCS_N2_ distributions (see Methods), as a proxy for an ensemble-averaged measurement of CCS (See Methods). Similar to the computationally derived R_g_ values, we observe an increase in CCS as a function of NaCl concentration in both AmAc and Tris-HCl infering that αS populates more extended conformations (Figure 4b,c). Examination of representative structures from the simulations at ionic strengths of either 10 mM or 150 mM demonstrates the role of interactions between the N- and C-termini of αS in mediating chain collapse (Figure 4d), consistent with previous reports^[34,95]^.

**Figure 4.**
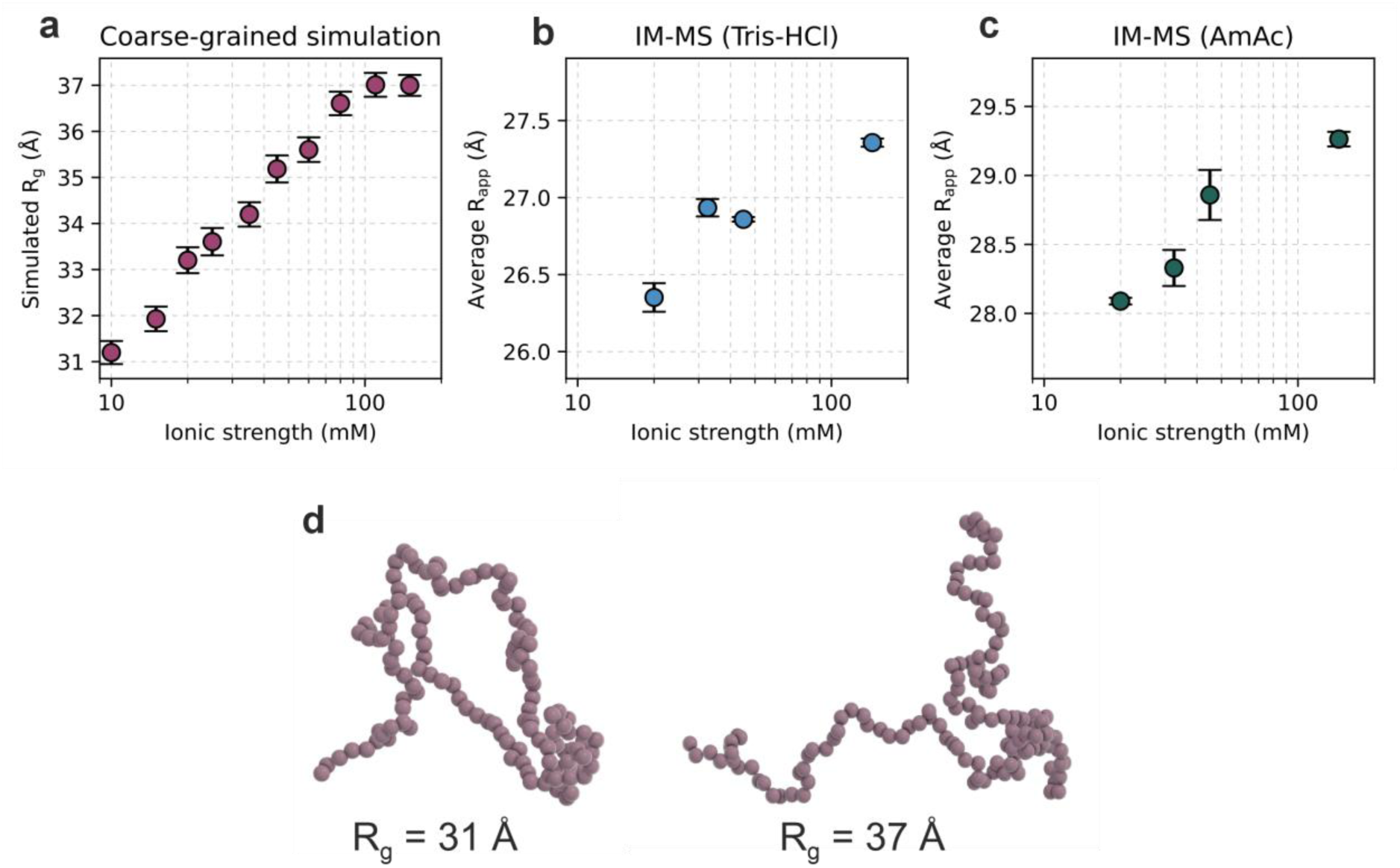
The mean R_g_ of αS from coarse grained simulations and average (R_app_) IM-MS values both increase in response to increasing ionic strength. (a) The mean radius of gyration (R_g_) of αS as determined by coarse-grained implicit-solvent simulations at varying ionic strengths (see Methods). The error shown is the standard error of the mean, determined by considering the calculated R_g_ values for all αS models in the simulation after a 5 ns equilibration (see Methods for details). (b, c) Average R_app_ calculated from the average ^TW^CCS_N2_ distributions of αS (see Methods, data from Figure 3) in (b) 20 mM Tris-HCl or (c) 20 mM AmAc, with data recorded in the presence of different concentrations of NaCl. (d) Representative coarse grained simulation frames of αS with calculated R_g_ values of 31Å and 37Å.

Next, we sought to compare our gas phase ^TW^CCS_N2_ measurements with solution phase measurements using diffusion-ordered spectroscopy nuclear magnetic resonance (DOSY NMR)^[96]^. We measured diffusion coefficients for αS in the absence or presence of NaCl (25 mM NaCl) in both AmAc and Tris-HCl buffers (Figure 5a,b,c and Figure S10). The data show that upon addition of NaCl, under both sets of buffer conditions, the measured diffusion coefficient of αS decreases (∼15%). The diffusion coefficient is inversely proportional to the hydrodynamic radius (R_h_) of the molecule, suggesting that addition of NaCl results in expansion of αS under the buffer conditions tested^[96,97]^. This is consistent with previous small angle X-ray scattering (SAXS) data that identified a NaCl-dependent increase in the R_g_ of αS^[19]^. Taken together, these data provide experimental evidence that IM-MS ^TW^CCS_N2_ gas-phase measurements are reflective of the solution-phase conformational distributions of αS, and demonstrates that it is possible, using nanopipette IM-MS, to determine the conformational properties of αS in different buffer conditions.

**Figure 5.**
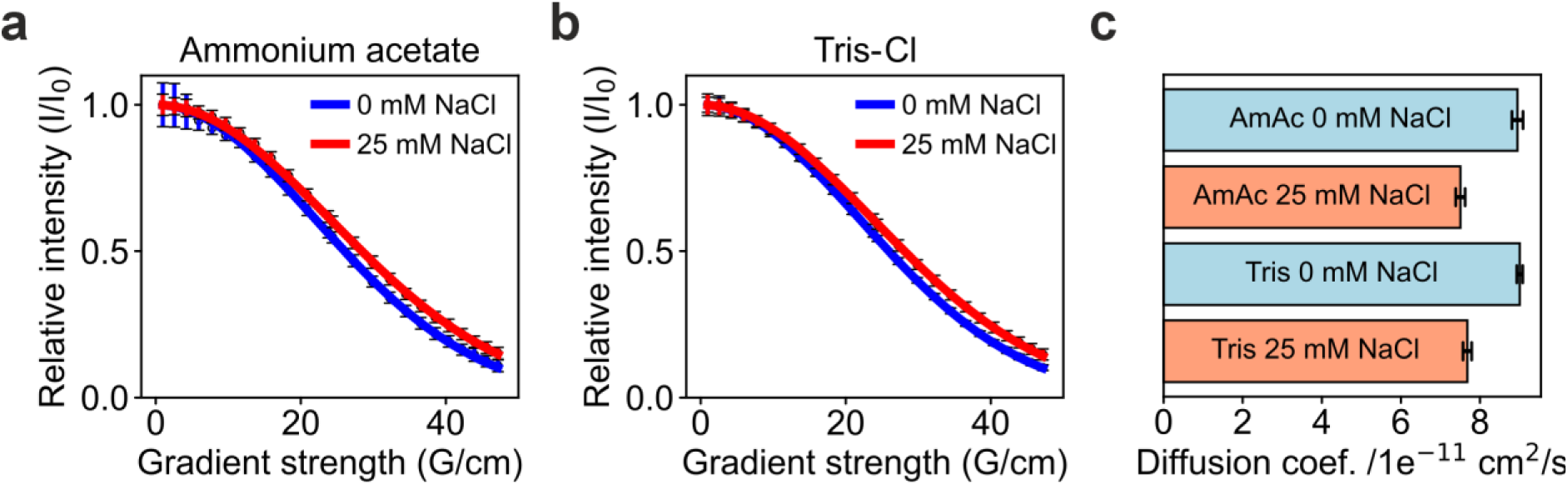
^1^H DOSY NMR indicates expansion of αS upon addition of NaCl. Diffusion coefficients for αS in solution are reduced in the presence of 25 mM NaCl. (a,b) Fitted ^1^H DOSY NMR data for αS in the presence or absence of 25 mM NaCl in (a) AmAc and (b) Tris-HCl buffer. Errors were calculated from the signal-to-noise level in the ^1^H spectra (see Methods). (c) αS diffusion coefficients under different buffer conditions. Errors were calculated using a Monte Carlo simulation approach and the spectral noise (see Methods). Samples contained 50 µM αS, 0 mM or 25 mM NaCl, in 100 mM ammonium acetate, pH 7.2, or 100 mM Tris-HCl, pH 7.2, at 24 °C.

To explore whether the conformational expansion of αS reflects a general sensitivity to ionic strength, or an effect on conformation that is specific to adding NaCl, we performed IM-MS of αS at elevated buffer concentrations (100 mM) of AmAc and Tris-HCl buffers. In the case of AmAc, at 20 mM the average CCS of the ^TW^CCS_N2_ distribution was 2487 Å^2^ (Figure 1b), and this increased to 2786 Å^2^ in 100 mM AmAc (Figure 6). In the presence of an additional 25 mM NaCl we observed a CCS shift of αS towards higher, more expanded ^TW^CCS_N2_ values, with the more compact conformational families significantly reducing in abundance which is also reflected in the measured charge state distributions (Figure 6, Figure S5). A similar shift was observed in Tris-HCl buffer upon addition of NaCl (Figure 6, S6). This suggests that the conformational changes that occur in elevated ionic strength conditions are not specific to Na^+^/Cl^−^ interactions but are instead driven by the overall ionic strength of the solution. To summarise, we find that R_app_ derived from IM-MS measurements in the gas phase, computationally calculated R_g_ values from CALVADOS simulations and diffusion coefficients by DOSY NMR all show a clear expansion/increase in size of αS upon addition of NaCl (Table S1).

**Figure 6.**
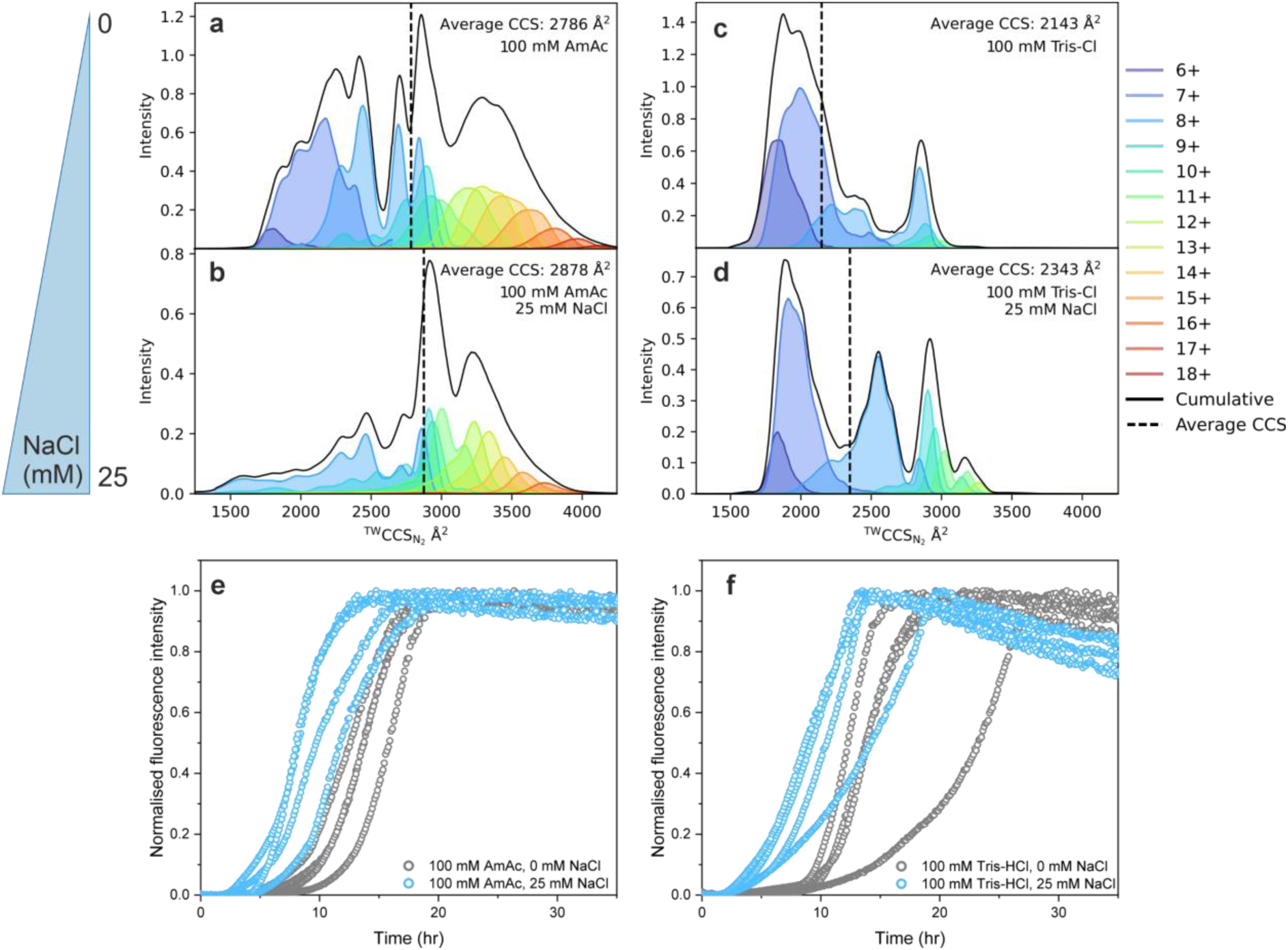
αS undergoes expansion at high ionic strength. ^TW^CCS_N2_ distributions showing the effect of NaCl on αS. αS was measured in (a) 100 mM AmAc with the addition of (b) 25 mM NaCl, and in (c) 100 mM Tris-HCl with the addition of (d) 25 mM NaCl. The key on the righthand side indicates the colour of the CCS distribution for each charge state, the black solid line is the cumulative fit and the black dotted linke represents the average ^TW^CCS_N2_ value. (e) ThT fluorescence in 100 mM AmAc in the absence or presence of 25 mM NaCl, and in (f) 100 mM Tris-HCl in the absence or presence of 25 mM NaCl. Three replicates are shown.

Environmental factors including salt concentration are known to determine amyloid fibril assembly conditions, leading to specific αS fibril morphologies^[98]^. Ions can stabilise both intra- and intermolecular hydrophobic interactions through dehydration, promoting fibril assembly^[20,99–101]^. Additionally, salt has been proposed to cause expansion of αS, exposing the hydrophobic amyloid prone NAC domain of the protein leading to accelerated amyloid assembly kinetics^[19]^. Here, in order to relate conformational changes to amyloid assembly kinetics were monitored using Thioflavin T (ThT) fluorescence in the presence or absence of NaCl (25 mM). The results showed that the rate of amyloid assembly increases with the addition of 25 mM NaCl compared to 0 mM NaCl in both AmAc and Tris-HCl buffers (Figure 6e,f)^[25,102]^. This suggests that the change in structure we observe with IM-MS directly impacts the amyloid assembly kinetics of αS, consistent with previous reports^[36]^. These findings together exemplify the role that structural rearrangements of αS measured by IM-MS correlate with its aggregation propensity and could help to elucidate disease relevant conformational families for therapeutic intervention.

## CONCLUSION

This study demonstrates that there are remarkable shifts in the conformational ensembles of αS driven by changes in the ionic environment, demonstrating that the structural landscape of αS is highly affected by salt concentration. The incorporation of nanopipette emitters has enabled us to show using native IM-MS that NaCl and the buffer composition modulate the conformational landscape of αS. We show that the conformational expansion of αS in presence of NaCl (Figure 3) determined using IM-MS correlates with in solution DOSY NMR measurements and computationally-derived R_g_ values (Figure 4, 5). IM-MS is well placed to capture snapshots of entire conformational ensembles and provide rich structural information for IDPs. Whereas a standard nESI set-up mostly limits analysis to volatile buffers which do not represent physiological conditions, nanopipette emitters enable new capabilities for IM-MS through the analysis of samples in common biochemical buffers and high-salt environments.

Overall, a clear reduction in αS salt adducts is observed in native nESI mass spectra using the nanopipette emitters descriped here (Figure 1, Figure 2 and Figure S3) which we attribute to the finer droplet spray^[103,104]^. The smaller droplets that emerge from the nanopipette partition the solvent into smaller volumes that contain fewer salt ions on average which reduces formation of adducts with proteins observed with standard ESI emitters. Our application of nanopipette emitters opens the door to deploying native IM-MS to afford a new understanding of how the conformations of IDPs relate to their function or dysfunction, such as the role of αS in synucleopathies, and to analyse proteins in buffers closer to their native cellular environment. Our results pave the way for the structural analysis of proteins under physiological conditions by native IM-MS. Future studies exploring the addition of other buffer components, and the introduction of complex sample environments such as cellular compartments and cell lysates, will further push this methodology forwards.

## ACKNOWLEDGEMENTS

A.N.C. and E.J.B. acknowledge support through a Sir Henry Dale Fellowship jointly funded by Wellcome and the Royal Society (220628/Z/20/Z). J.A.C acknowledges the support of the University of Leeds and the BBSRC (BB/Y00034X/1). P.A., C.C.C.C. and A.N.C. acknowledge funding from the BBSRC (BB/X003086/1) and F.S., P.A., E.L.N. from the MRC (MR/W031515/1). S.E.R. is the grateful recipient of a Royal Society Research Professorship (RSRP/R1/211057). Funding from the BBSRC enabled the purchase of MS equipment (BB/E012558/1). The authors thank Dr Sri Ranjani Ganji and Dr Gemma Wildsmith from the Biomolecular Mass Spectrometry Facility at the University of Leeds for their support and assistance in this work. For access to the 600 MHz spectrometer, we acknowledge the Astbury Biostructure Laboratory BioNMR facility which was funded by the University of Leeds. We also thank Arnout Kalverda for excellent technical support, and Dr Pijush Chakraborty and Dr Alex Heyam for helpful discussions and advice.

## METHODS

### Standard nESI emitter fabrication

Borosilicate thin wall glass capillaries (0.78 mm) with filament (Harvard Apparatus, UK) were pulled using a P-97 Flaming/Brown micropipette puller (Sputter Instrument, CA, USA) to make two nanospray tips. The optimised procedure used three cycles as follows; heat 560, pull 250, velocity 10, time 50 s. The pulling protocol is specific to the instrument and can vary between different pullers and filaments. Capillaries were placed into a glass petri dish and coated with palladium in a SC7620 mini sputter coater (Quorum Technologies, Sussex, UK) pressurised with argon at 2 x 10^−2^ mbar. When a current of 25-30 mA was applied the plasma enabled palladium atoms to deposit on the glass surface. The coating time lasted 75 s after which the petri dish was rotated by 90° followed by a second coating. Samples (10 µL) were loaded into capillaries which were then clipped to size using tweezers.

### Nanopipette nESI emitter fabrication

The nanopipette nESI emitter tips were fabricated using 1.0 mm outer diameter and 0.5 mm inner diameter quartz capillaries (QF100-50-7.5; Sutter Instrument) with the SU-P2000 laser puller (World Precision Instruments). A two-line protocol was used: line 1 with HEAT 750/FIL 4/VEL 30/DEL 150/PUL 80, followed by line 2 with HEAT 850/FIL 3/VEL 40/DEL 135/PUL 225. The pulling protocol is specific to the instrument and can vary between different pullers. For native IM-MS measurements, the emitters were loaded with analyte solution (8 µL) and fitted with a platinum wire (PT00-WR-000117; Goodfellow) prior to use.

### Scanning electron microscopy (SEM) nESI emitter characterisation

The pore dimensions of the standard nESI and nanopipette nESI were characterised by field emission SEM with a FEI Nova 450 at an accelerating voltage of 3–5 kV. Images were prepared without coating, using a CBS detector (back-scattered electron detector).

### Native ion mobility mass spectrometry

Native IM-MS experiments were performed on a Waters Synapt HDMS mass spectrometer with travelling (T-wave) ion mobility and a nano-ESI source. N-terminally acetylated αS, expressed and purified as described in^[36]^, was analysed at a concentration of 20 µM in 20 mM AmAc, 20 mM Tris-HCl, 1 x PBS (Dulbeccos A; Thermo Fisher Scientific), 10 mM potassium phosphate, 20 mM sodium phosphate, 100 mM AmAc, and 100 mM Tris-HCl each at a pH of 7.2. For salt titrations, NaCl was added at 12.5 mM, 25 mM and 125 mM. Instrument parameters were set at: capillary voltage 1.0 kV, source temperature 30 °C, backing pressure 0.0 – 0.3 Bar, sampling cone 18 V, extraction cone 1.0 V, trap collision energy 5 V, transfer collision energy 2.0 V, trap DC bias 30 V, IM wave velocity 300 m/s, IM wave height 7.0 V. Gas pressures in the instrument were: trap cell 0.0258 mbar, IM cell 0.36 mbar. ^TW^CCS_N2_ signifies CCS values calculated using traveling wave ion mobility in N_2_ buffer gas using calibrants acquired in N_2_ buffer gas. Lysozyme was analysed at 10 µM under identical conditions except for: sampling cone 10 V and trap DC bias 12 V. MassLynx V4.1 (Waters Corporation) was used for data processing. The IM spectra were calibrated according to the Bush database^[46]^ using denatured cytochrome C (charge states 13+ to 19+), myoglobin (charge states 15+ to 24+) and ubiquitin (charge states 7+ to 13+) at 10 µM in 50 % (v/v) acetonitrile, 0.1 % (v/v) formic acid. Calibrated intensity values were normalised using a scaling factor determined by the signal intensity in the mass spectrum as well as the area under the arrival time distribution peak. Hen egg white lysozyme (Sigma) IM spectra were also calibrated^[46]^ using native cytochrome C (charge states 6+ and 7+) and myoglobin (charge states 7+ to 9+) measured in 100 mM AmAc. The apparent radius of gyration (R_app_) was calculated using the following equation: 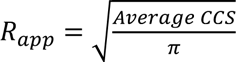

### ^1^H DOSY NMR

DOSY experiments were performed on a 600 MHz Bruker Avance III spectrometer equipped with a 5 mm QCI-P CryoProbe. For each DOSY experiment under each sample condition, a total of 24 1D acquisitions were performed at different gradient strengths between 2% and 98% where the maximum gradient strength was 48.148 G cm^−1^. To improve data fitting the gradient strength percentages were weighted quadratically towards both 2% and 98% as follows: 2.00, 5.32, 8.76, 12.30, 16.10, 20.00, 24.10, 28.40, 33.00, 38.00, 43.40, 49.40, 50.60, 56.60, 62.00, 67.00, 71.60, 75.90, 80.00, 83.90, 87.70, 91.20, 94.70, 98.00. Following processing, peaks were integrated in the methyl region between 0.6-0.9 ppm. To obtain the error for each experiment, the noise at each gradient strength was obtained by integrating a region of the baseline of the same size where no peaks were present (0.3-0.6 ppm). To obtain the diffusion coefficient, following baseline correction and integration, the data were fitted to the following equation:

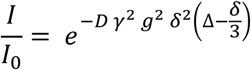

where *I* is the observed intensity, *I_0_* the reference intensity (the signal intensity of the first point), *D* is the diffusion coefficient, γ is the gyromagnetic ratio of the observed nucleus, *g* is the gradient strength, δ the length of the gradient, and Δ the diffusion time. The length of the gradient (δ) and diffusion time (Δ) were 2 ms and 400 ms, respectively. Baseline adjustment, integration, fitting and plotting was performed with in-house Python3 scripts which made use of the NumPy (v1.26.4)^[100]^, SciPy (v1.13.1)^[101]^ and Matplotlib (v3.9.2)^[102]^ libraries. The integration was performed using the numpy.trapz function. To estimate the uncertainty in the diffusion coefficient a Monte Carlo simulation approach, incorporating the errors associated with each data point, was employed. To simulate the effect of the error in each data point on the diffusion coefficient, we first generated 1000 synthetic datasets. In each dataset, every original integrated signal intensity was perturbed by adding random noise drawn from a Gaussian distribution with a mean of zero and a standard deviation equal to the error of that data point. To determine the diffusion coefficient and associated error for each experiment we used the mean and standard deviation of fits to these 1000 datasets. Samples contained 50 µM α-synuclein, 0 mM or 25 mM NaCl, in 100 mM ammonium acetate, pH 7.2, or 100 mM Tris-HCl, pH 7.2, at 24 °C.

### ThT amyloid assembly kinetics

Kinetics of αS amyloid formation were monitored in a 96-well, non-binding, flat-bottom, half-area microplate (Corning, USA; 10438082) containing one Teflon polyball (1/8” diameter; Polysciences Europe) per each well of sample. Samples of 100 µL containing 100 µM aS with 20 µM Thioflavin T (ThT) in 100 mM ammonium acetate, pH 7.2 and 100 mM Tris-HCl with 0 mM NaCl and 25 mM NaCl were incubated at 37 °C shaking at 600 rpm in a FLUOstar omega plate reader (BMG Labtech). Fluorescence intensity was measured by exciting at 440 ± 10 nm and collecting emission at 482 ± 12 nm using a bandpass filter. Results were blank corrected and normalised to the maximum fluorescence value of each curve.

### Coarse-grained simulations

Coarse-grained implicit-solvent simulations of αS were performed in the in the OpenMM framework (version 8.1.1)^[108]^ using the CALVADOS python package and the CALVADOS2 force field^[93,94]^. Unless otherwise stated, simulation parameters used were kept as their default values in the software. The N terminal positive charge was removed to match the N-terminally acetylated sample used in the MS and NMR experiments. Simulations were performed in a 30 nm^3^ box at pH 7.5. Following energy minimization, simulations were run at a temperature of 25 °C. Simulations were performed for a total of 500 ns each with coordinates being saved every 0.5 ns. The simulation had a time step of 10 fs and a friction coefficient of 10 fs^−1^. The first 5 ns of each simulation was considered to be part of the equilibration time and therefore discarded. In total, 10 simulations were carried out at ionic strengths of 10, 15, 20, 25, 35, 45, 60, 80, 110 and 150 mM. The R_g_ was calculated using the inbuilt function in the CALVADOS software for each of the 990 frames outside the equilibration time, with the error bars representing the standard error of the mean.

