## Supplemental Figures for "Nanopipettes enable native electrospray ionisation ion mobility mass spectrometry studies of buffer-dependent conformational dynamics of α-synuclein"

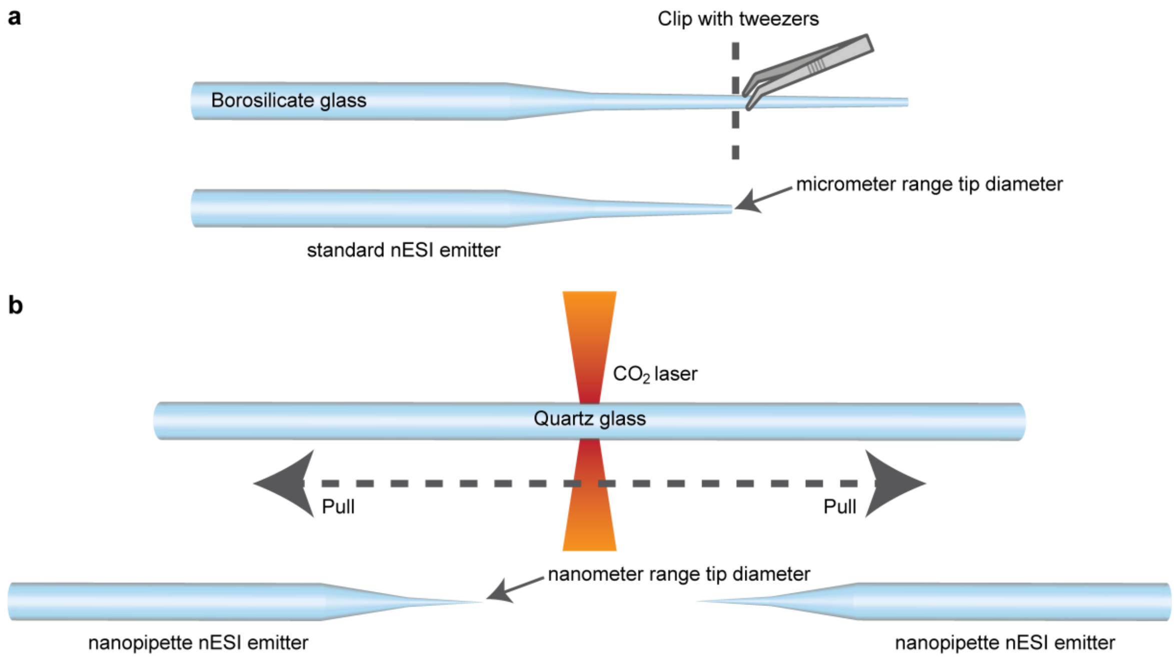


***Figure S1. Fabrication of the nESI emitter tips.*** *The typical fabrication process for the standard nESI emitter tip (a) starts with a pre-pulled borosilicate glass pipette with a long taper. The long taper is clipped with tweezers, resulting in a tip diameter in the 2 - 20 micrometre range. For the fabrication of the nanopipette nESI emitter tip (b), a laser puller is used with quartz glass capillaries. This process results in the fabrication of a pair of nearly identical nanopipette nESI emitter tips, each with a nanometre range sized pore.*

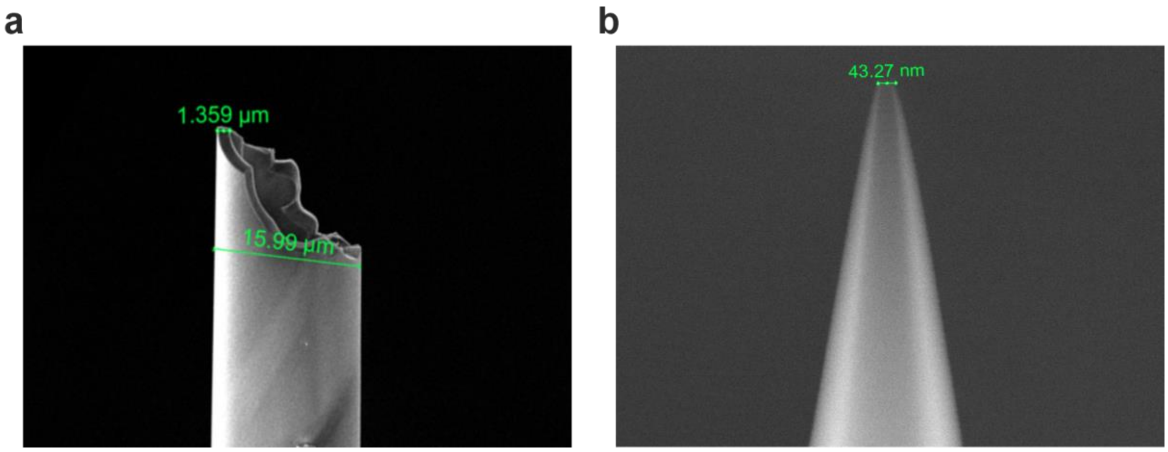


***Figure S2. Comparison of nanopipette emitters with standard nESI emitters****. SEM images of (a) standard borosilicate nESI emitters clipped to size with tweezers (pore diameter ~ 16 µm) and (b) of nanopipette emitters (pore diameter ~ 43 nm).*

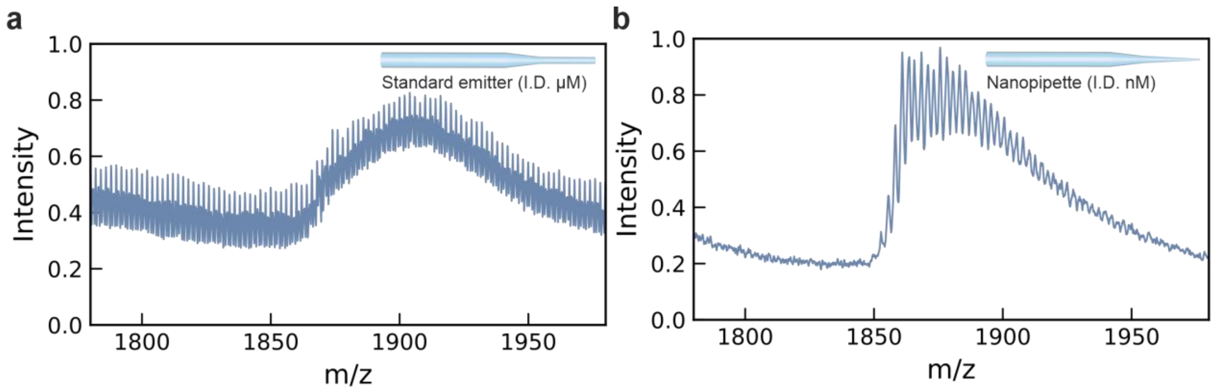


***Figure S3. Zoomed in native nESI mass spectra of the 8+ charge state of αS.*** *Measured in PBS, pH 7.2 using (a) standard borosilicate emitters and (b) quartz nanopipettes. The predicted m/z of the apo 8+ αS charge state is 1813. The full mass spectra are shown in Figure 1.*

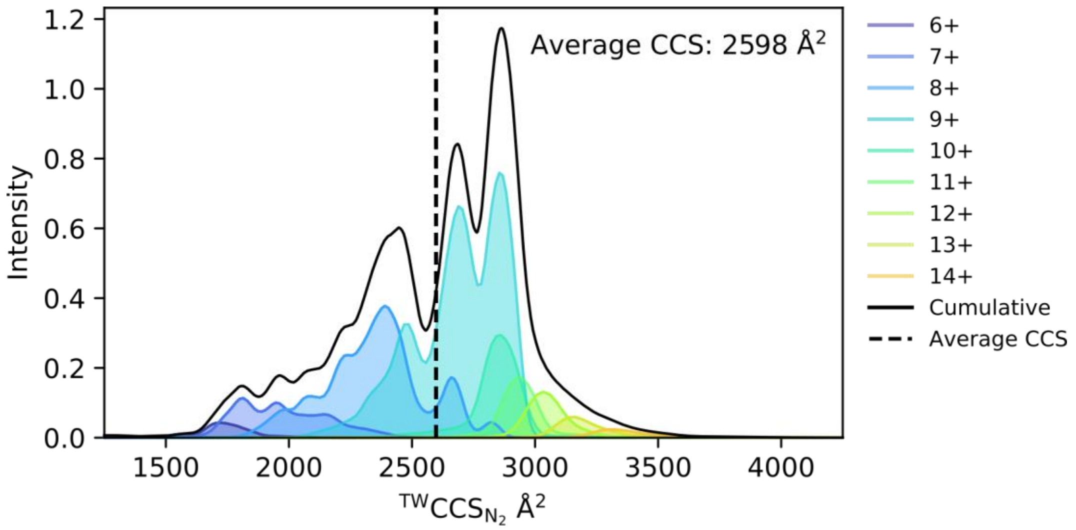


***Figure S4. Representative $^{TW}CCS_{N2}$ distribution of αS in PBS buffer using a nanopipette nESI emitter.*** *The data are taken from a spectrum acquired of 10 µM αS in PBS buffer, pH 7.2. The key on the righthand side indicates the colour of the CCS distribution for each charge state, the black solid line is the cumulative fit and the black dotted line represents the average $^{TW}CCS_{N2}$ value.*

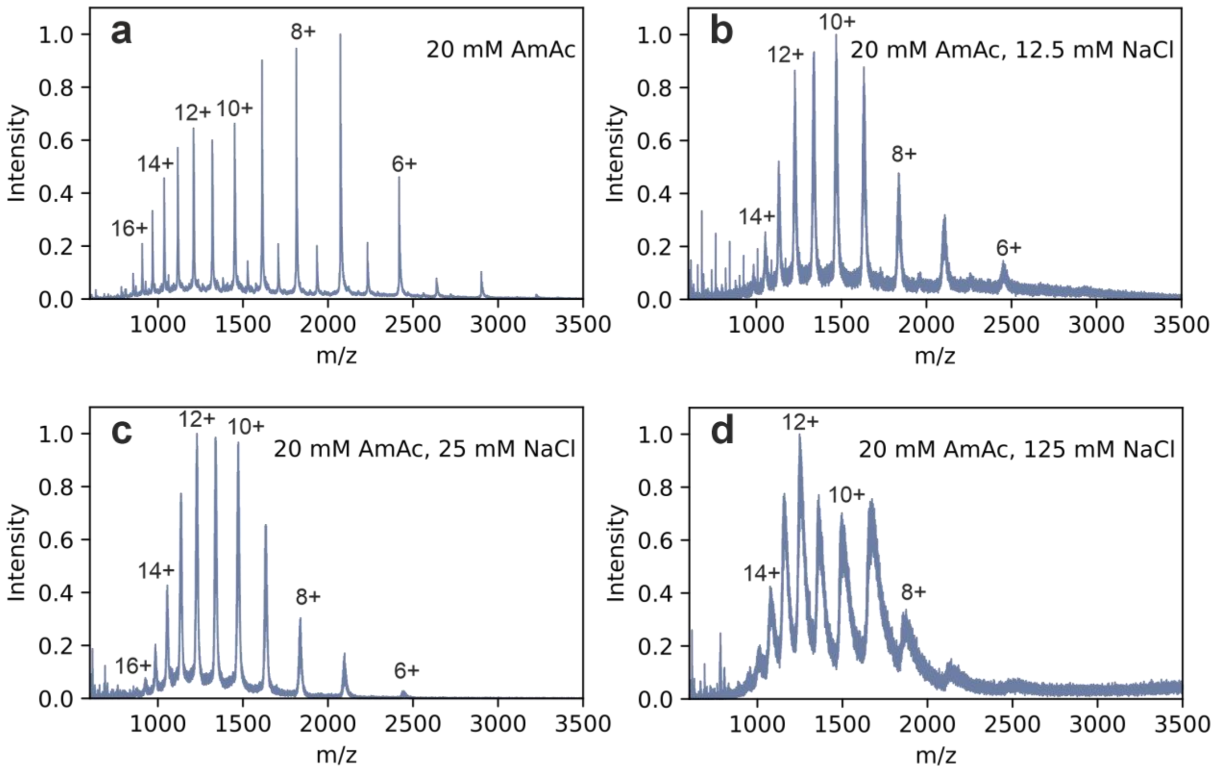


***Figure S5. Native nESI mass spectra of αS in 20 mM AmAc with NaCl titration.*** *αS (20 µM) was measured in (a) 20 mM AmAc, pH 7.2 with the addition of (b) 12.5 mM NaCl, (c) 25 mM NaCl, (d) 125 mM NaCl.*

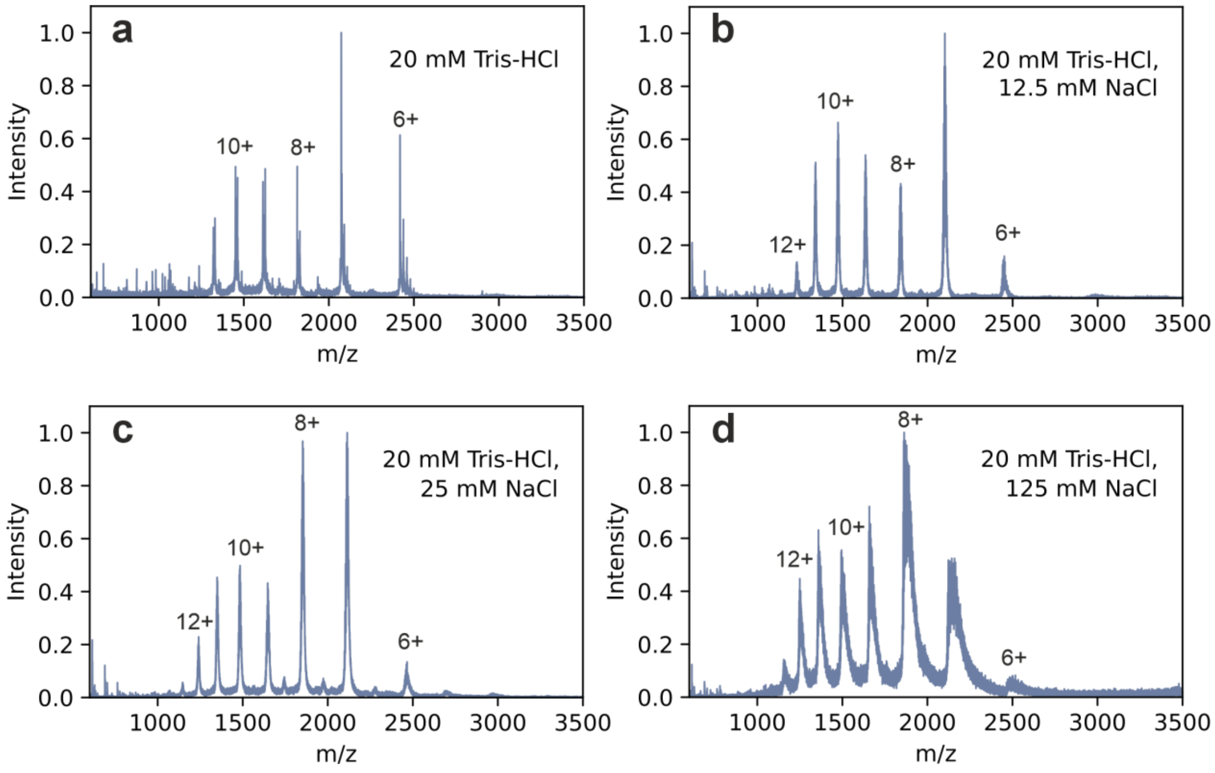


***Figure S6. Native nESI mass spectra of αS in 20 mM Tris-HCl with NaCl titration.*** *αS (20 µM) was measured in (a) 20 mM Tris-HCl, pH 7.2 with the addition of (b) 12.5 mM NaCl, (c) 25 mM NaCl, (d) 125 mM NaCl.*

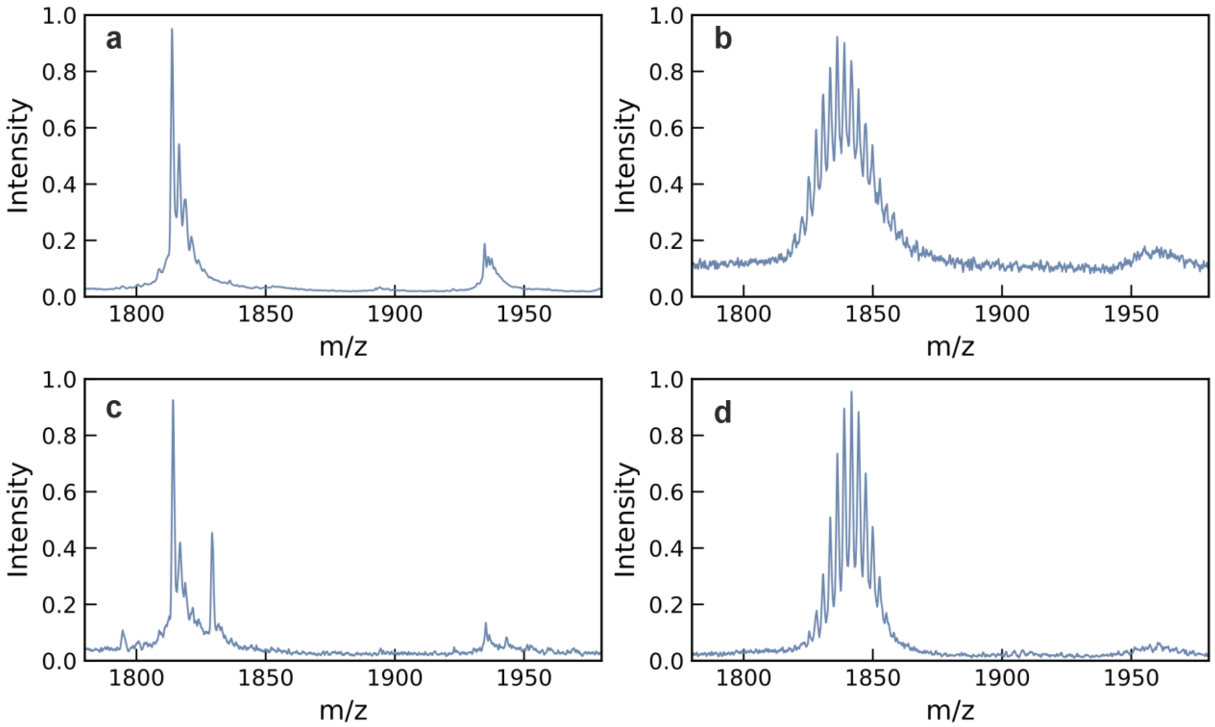


***Figure S7. Native nESI mass spectra of the 8+ charge state.*** *αS measured in (a) 20 mM AmAc, (b) 20 mM AmAc with 12.5 mM NaCl, (c) 20 mM Tris-HCl and (d) 20 mM Tris-HCl with 12.5 mM NaCl. All measurements were taken at pH 7.2.*

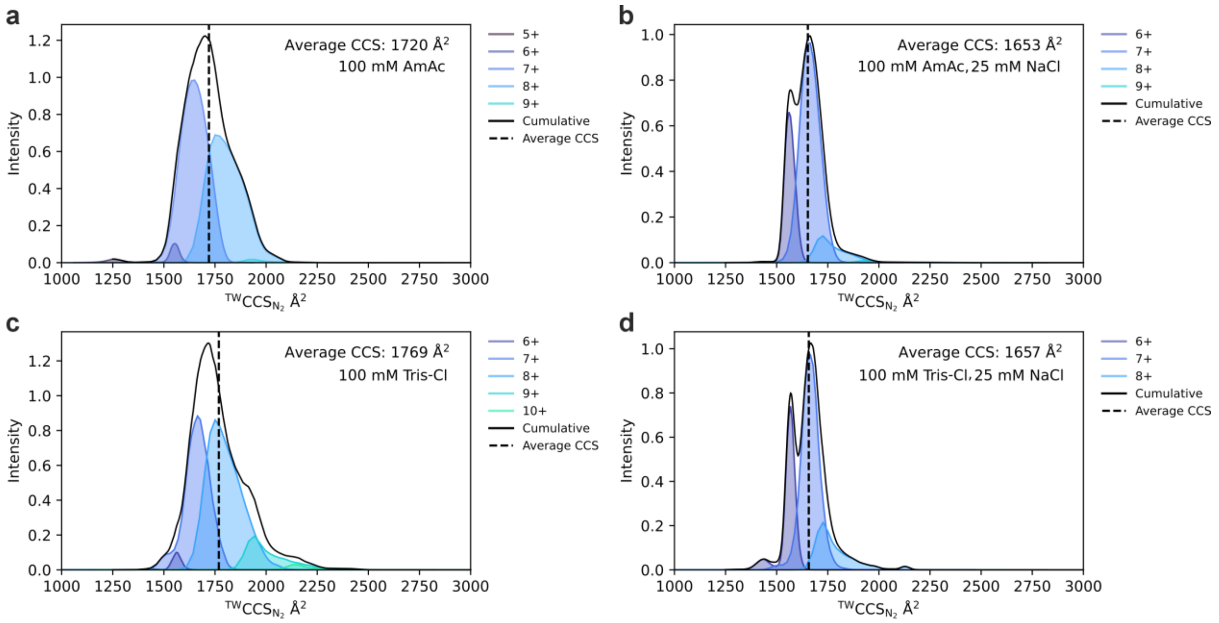


***Figure S8.*** $^{TW}$***CCS***$_{N2}$ ***Ion mobility distributions of lysozyme using nanopipette nESI emitters.*** *nESI mass spectra were acquired of 10 µM hen egg white lysozyme in (a) 100 mM AmAc, (b) 100 mM AmAc, pH 7.2 with 25 mM NaCl, (c) 100 mM Tris-HCl and (d) 100 mM Tris with 25 mM NaCl.*

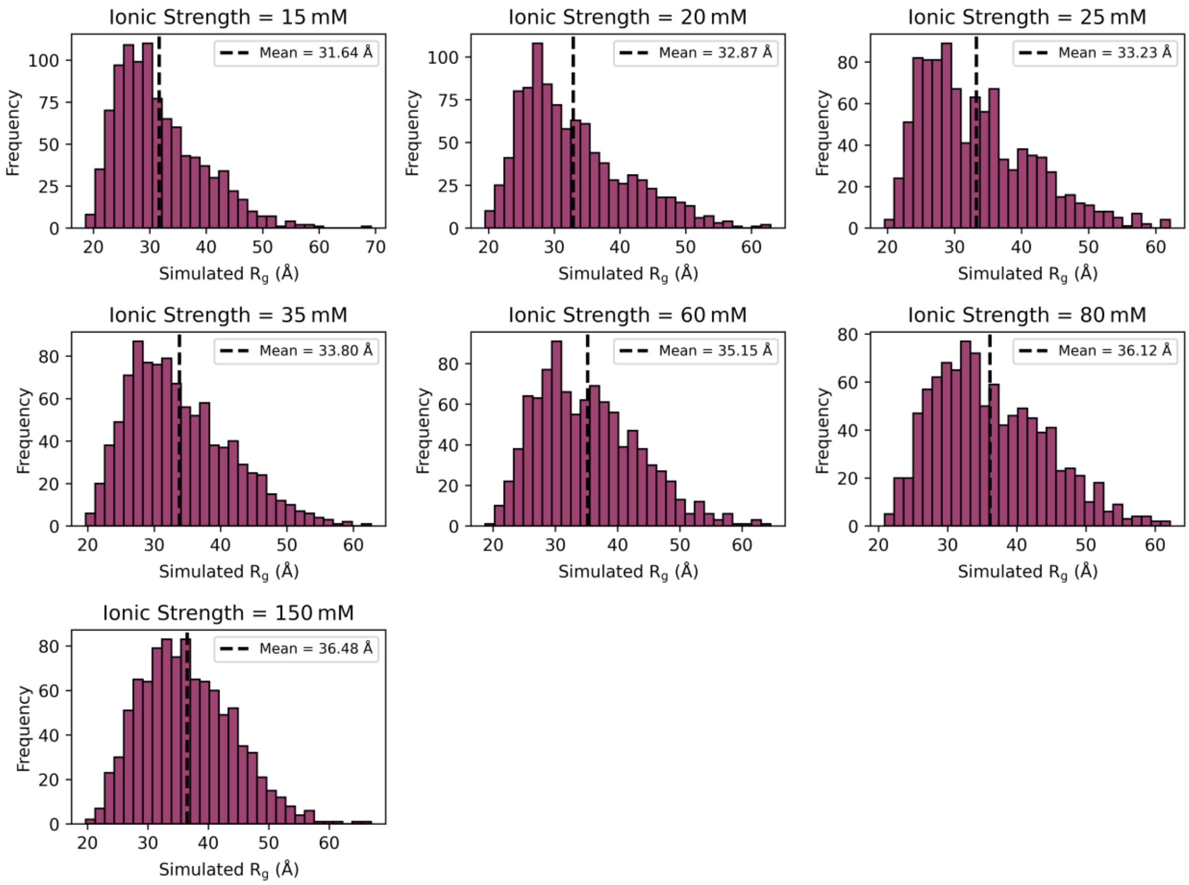


***Figure S9. Distributions of calculated Rg values for αS from coarse grained molecular dynamics simulations.*** *Each panel shows a different ionic strength value for the simulation. The mean* $R_g$ *is shown as a vertical black dashed line. For details on the simulation and number of frames used, see Methods.*

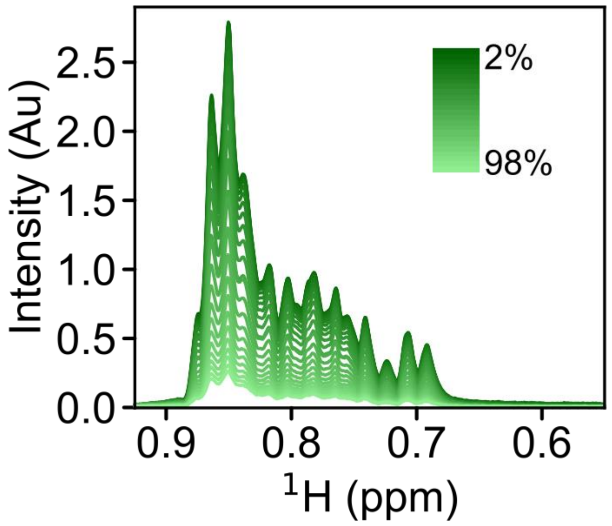


***Figure S10. Example 1D spectra of αS from DOSY NMR experiments.*** *For each experiment under each sample condition, a total of 24 1D acquisitions were made at different gradient strengths between 2% and 98%, where the maximum gradient strength was 48.148 G cm*$^{-1}$*. Data for the region used for integration (0.6-0.9 ppm) are shown. The sample contained 50 µM αS in 100 mM Tris-HCl, pH 7.2, at 24 ˚C.*

| Ionic Strength | $R_{app}$ (Å) | % change | $R_g$ | % change | Diffusion coefficient × $10^{-7}$ ($cm^2$ $s^{-1}$) | % change |
|---|---|---|---|---|---|---|
| 20 mM | 28.14 | +8% | 32.87 | +11% | 8.99 ± 0.05 | -16.2% |
| 125 mM | 30.27 | | 36.48 | | 7.53 ± 0.04 | |

***Table S1. Table comparing measurements of αS expansion at 20 mM and 125 mM ionic strength.*** *$R_{app}$ in Å was calculated from the average $^{TW}CCS_{N2}$ values for αS in 20 mM AmAc, pH 7.2 and 100 mM AmAc with 25 mM NaCl, pH 7.2 (see methods for calculation). $R_g$ was calculated from coarse grained CALVADOS simulations of αS at 20 mM and 125 mM ionic strength, pH 7.5. Diffusion coefficient ($cm^2$/s) were measured using DOSY NMR.*
